# Complex small-world regulatory networks emerge from the 3D organisation of the human genome

**DOI:** 10.1101/2020.05.12.091041

**Authors:** C. A. Brackley, N. Gilbert, D. Michieletto, A. Papantonis, M. C. F. Pereira, P. R. Cook, D. Marenduzzo

## Abstract

The discovery that overexpressing one or a few critical transcription factors can switch cell state suggests that gene regulatory networks are relatively simple. In contrast, genome-wide association studies (GWAS) point to complex phenotypes being determined by hundreds of loci that rarely encode transcription factors and which individually have small effects. Here, we use computer simulations and a simple fitting-free polymer model of chromosomes to show that spatial correlations arising from 3D genome organisation naturally lead to stochastic and bursty transcription plus complex small-world regulatory networks (where the transcriptional activity of each genomic region subtly affects almost all others). These effects require factors to be present at sub-saturating levels; increasing levels dramatically simplifies networks as more transcription units are pressed into use. Consequently, results from GWAS can be reconciled with those involving overexpression. We apply this pan-genomic model to predict patterns of transcriptional activity in whole human chromosomes, and, as an example, the effects of the deletion causing the diGeorge syndrome.

## INTRODUCTION

Transcription – the copying of DNA into RNA – is tightly regulated. Early insights into regulatory mechanisms came from work on binary on/off genetic switches controlled by one (or just a few) transcription factors such as the lambda and lac repressor in *Escherichia coli* [1]. Similar regulatory mechanisms are present in eukaryotes, albeit with additional complexity. For instance, a fibroblast cell can be reprogrammed into a muscle cell by a single master regulator (MYOD) [2, 3], or into pluripotent stem cells by four Yamanaka factors (Oct4, Sox2, c-Myc, Klf4) [4].

Genome-wide association studies (GWAS) lead to quite a different view: gene regulation is widely distributed and involves interactions between hundreds (perhaps thousands) of loci scattered around the genome [5, 6]. GWAS allows quantitative trait loci (QTLs) affecting any measurable genetic trait to be ranked in an unbiased way. With complex traits like human height, and diseases such as schizophrenia and type II diabetes, the top ten QTLs in the rank order combine to yield only modest effects, while the top one-hundred still account for less than half of the total genetic effect. Hundreds more QTLs are expected to be identified as sample sizes and data resolution improve [5–7]. Expression QTLs (eQTLs) are QTLs affecting transcription of other DNA regions. Perhaps surprisingly, these are rarely found in genes encoding transcription factors or other proteins; instead, they usually involve single-nucleotide changes in non-coding elements that bind transcription factors such as active enhancers and promoters [8–10].

Results from GWAS lead to the view that most gene-regulatory networks are incredibly complex, with the activity of a given gene being affected by a panoply of eQTLs, each having only a tiny effect. This is captured by the “omnigenic” model, which is based on a set of gene-interaction equations [5, 6] such that the activity of almost any gene affects that of almost every other one. While this provides an excellent and appealing framework for viewing GWAS results, it requires many types of transcription factors to be biochemical mediators of interactions; therefore, it has limited predictive power due to the large number of free parameters. Moreover, it is openly short on molecular detail.

Here we propose an alternative framework that links transcriptional regulation with 3D genome structure. We explore this framework using stochastic computer simulations of a polymer model for chromosome organization, in which a chain of beads represents a chromatin fibre, and a set of spheres represent complexes of transcription factors and RNA polymerases – which we will call “TFs” for short. Some chromatin beads are identified as transcription units (TUs), and we shall call them TU beads. They contain binding sites for TFs, and can be sites of transcriptional initiation (we do not discriminate between genic and non-genic promoters and enhancers). As a simple starting point we only consider one type of TF that can bind specifically and multivalently to TU beads, and non-specifically (i.e., with weak affinity) to every other bead. We perform 3D Brownian dynamics simulations that evolve the diffusive dynamics of the chain and associated factors. We previously showed that similar polymer models can yield structures resembling those seen using chromosome-conformation-capture (3C) [11–15] and microscopy [16]. In this work, we link 3D structure to expression and transcriptional dynamics by measuring how often a TU bead is transcribed – which we do by computing the fraction of time over which a TF is bound to it. To establish the methodology, we model a short chain of beads each representing 3 Mbp of chromatin, before going on to simulate whole human chromosomes.

*In vivo*, one expects biochemical regulation to operate alongside any structural regulation. However, to enable unambiguous interpretation, we deliberately exclude specific biochemical regulation (e.g., we do not include multiple TF species, each binding to a subset of TUs, with TU activity feeding back on TF abundance). Our simulations nevertheless capture many features of eukaryotic regulation. For example, transcription is stochastic and bursty (in agreement with single-cell transcriptomics data), and small-world (percolating) networks that encapsulate much of the rich complexity observed in GWAS emerge through spatial effects alone. The resulting picture is similar to that given by the omnigenic model [5, 6], but the regulatory networks involve much more than the “core” and “peripheral” genes of that model; they additionally include non-genic TUs, chromatin loops, active euchromatin, and even inert heterochromatin. Consequently, the activity (or inactivity) of most (probably all) TUs in our model is affected by the activity of most (probably all) other segments in the genome. We find such pan-genomic regulation critically requires non-saturating concentrations of TFs – as normally found *in vivo* – and that increasing concentrations dramatically simplify networks. This enables us to reconcile the GWAS-based view that regulatory networks are highly complicated with the observation that overexpressing one or a few TFs can decisively alter cell state.

## RESULTS

We start by considering a simple system where a 3 Mbp chromatin fragment is represented by a chain of 1000 beads (each 30 nm in diameter, and corresponding to 3 kbp). We select at random *N* = 39 of these beads and identify them as TUs (Fig. 1A; full details in SI). Additionally, *n* spheres (also 30 nm in diameter) represent complexes of transcription factors and RNA polymerase II (called TFs for short). TFs bind reversibly to TUs via a strong attractive interaction, and to all other beads weakly and non-specifically. An important feature is that TFs switch between an active (binding) and an inactive (non-binding) state at rate *α*. Many factors switch like this *in vivo* (e.g., due to phosphorylation and de-phosphorylation), and switching is required to account for the rapid exchange of factors and polymerases between bound and free states seen in live-cell photobleaching experiments [17]. As ~ 7 out of 8 polymerases attempting to initiate at promoters also dissociate with a half-life of ~ 2.4 s [18], our complexes generally behave like those *in vivo*. We say a TU bead is transcribed whenever a TF lies close to it (Methods); the transcriptional activity of a TU is then the fraction of time it is transcribed during a simulation. To reflect the situation in mammalian cells [19], we initially assume there are fewer TFs than TU beads (i.e., *n* = 10 TFs in the active binding state at any time, compared to 39 TUs).

**FIG. 1:**
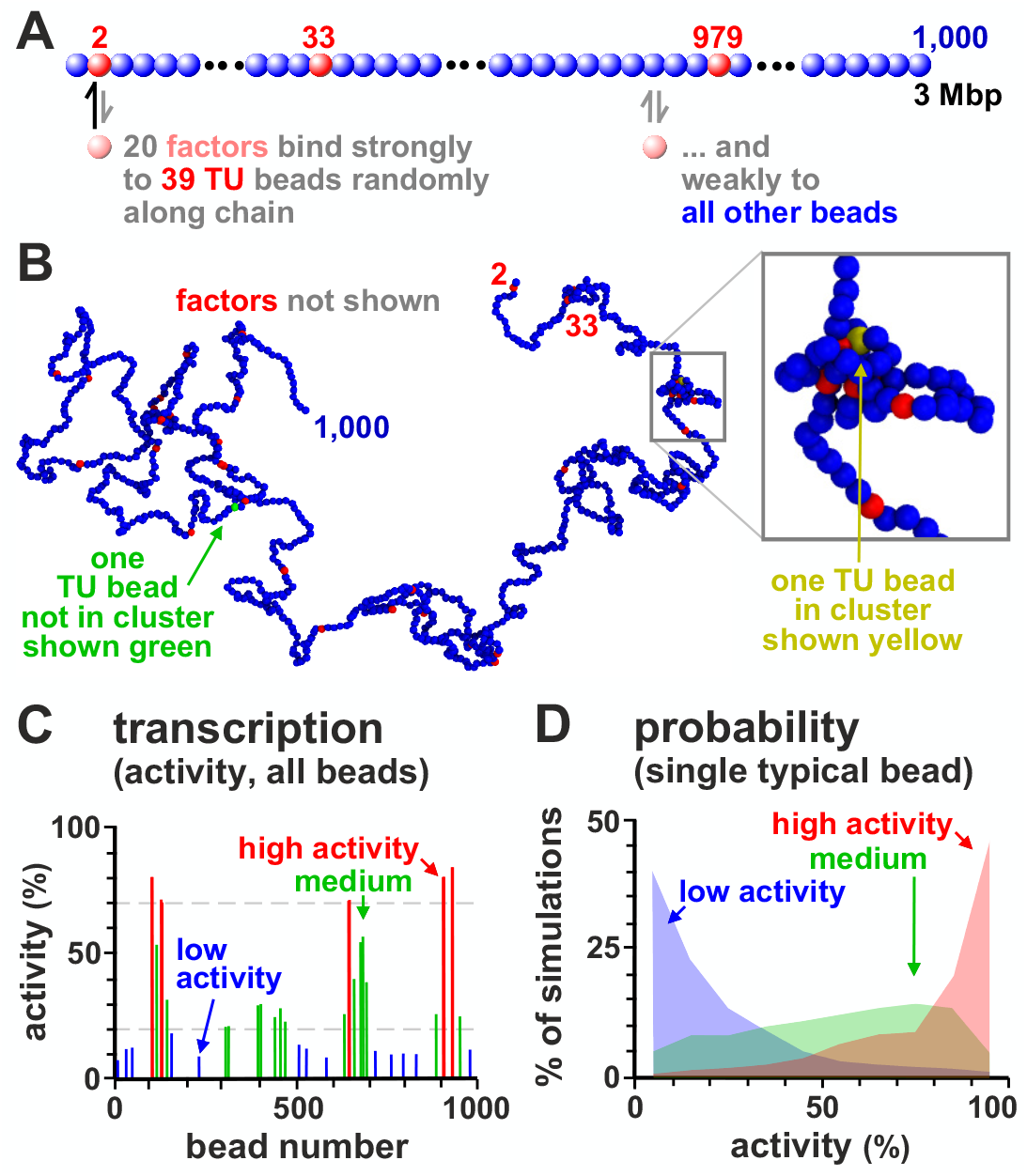
Patterns of transcriptional activity. **A.** Schematic of the model. Twenty TFs (pink) that switch between on/off states at rate 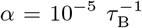 (with *τ*_B_ the Brownian time, see Methods) or 0.001 s^−1^ bind specifically to 39 TUs (red beads) randomly positioned along the chain, and non-specifically to the other beads (blue). A TU is considered transcriptionally active if associated with a TF. **B.** Example conformation (TFs not shown). Some beads cluster and form loops; one TU not in a cluster (and not transcribed) is green, and another that is in a cluster (and transcribed) is yellow. Inset: zoom of boxed region. **C.** Transcriptional activity for each TU bead averaged over 1000 simulations (each lasting 10^5^ *τ*_B_). TUs are grouped according to activity, with red, green and blue bars showing high (> 70%), medium (20 70%) and low (< 20%) activity, respectively. This gives a population-level measure of activity. **D.** Variation of activity across simulations (reflecting cell-to-cell variation) for 3 representative TUs with high (red), medium (green), or low (blue) average activity (defined as in **C**).

As noted above, our Brownian dynamics simulations allow us to predict transcriptional activity of TUs. By interrogating TF-chromatin interactions at regular time intervals over many repeat simulations we build up a population picture of transcription. A typical configuration of the 3 Mbp fragment is shown in Figure 1B. Strikingly, bound TFs spontaneously cluster, despite there being no attractive interactions between TUs or between TFs. Such clustering is driven by the “bridging-induced attraction” [12, 20, 21] that arises due to a positive feedback: when a TF forms a molecular bridge between two chromatin regions and forms a loop, the local chromatin concentration increases, making further TF binding more likely. Clusters then grow until limited by entropic costs of crowding (Fig. S1A). Most of the non-trivial phenomena described below result from such clustering, which in turn requires TF multivalency [20]. These clusters closely mirror those seen *in vivo*, which are variously described as transcriptional compartments, hubs, super-enhancer (SE) clusters, phase-separated droplets/condensates, and factories [7, 10, 22–24].

### Transcriptional activity varies along the chromatin fibre and is highly stochastic

As our model TFs have the same affinity for all TUs, one might expect each TU to be bound with equal likelihood; however, transcriptional activity (the fraction of time a TU is transcribed) varies from ~10–90% (Fig. 1C). What causes this variation? As TF copy number is limiting, and as bound TFs cluster, most transcription occurs in clusters (as is the case *in vivo* [7, 22]). Since TUs are positioned irregularly along the fragment, some have closer neighbours in 1D sequence space than others, and these are inevitably the ones most likely to cluster and be transcribed (Fig. S1B).

Whilst Figure 1C pertains to population averages of 1000 independent simulations, it is informative to consider each simulation independently (as in single-cell transcriptomics). Such analysis shows that transcriptional activity is stochastic, varying substantially from simulation to simulation: a TU which is on average highly-active may be silent in some simulations, while a TU which is relatively silent overall may be active in others (Fig. 1D).

### Transcriptional bursting

During a simulation, chromatin conformation can change dramatically (Fig. 2A). Such changes often yield transcriptional “bursts” – periods of continued activity followed by silent periods (Fig. 2B) – as TUs with intermediate levels of activity repeatedly join a cluster to give a burst and then dissociate. Notably, TUs lying close to each other in sequence space often start and stop bursts coordinately due to the intrinsic positive feedback in the system (Fig. S1A). These results are consistent with experimental observations: single cell Hi-C [25] and transcriptomics [26] show that the structure and function of each individual cell is unique, and bursting is well documented [27–30] with nearby promoters often firing together [28].

**FIG. 2:**
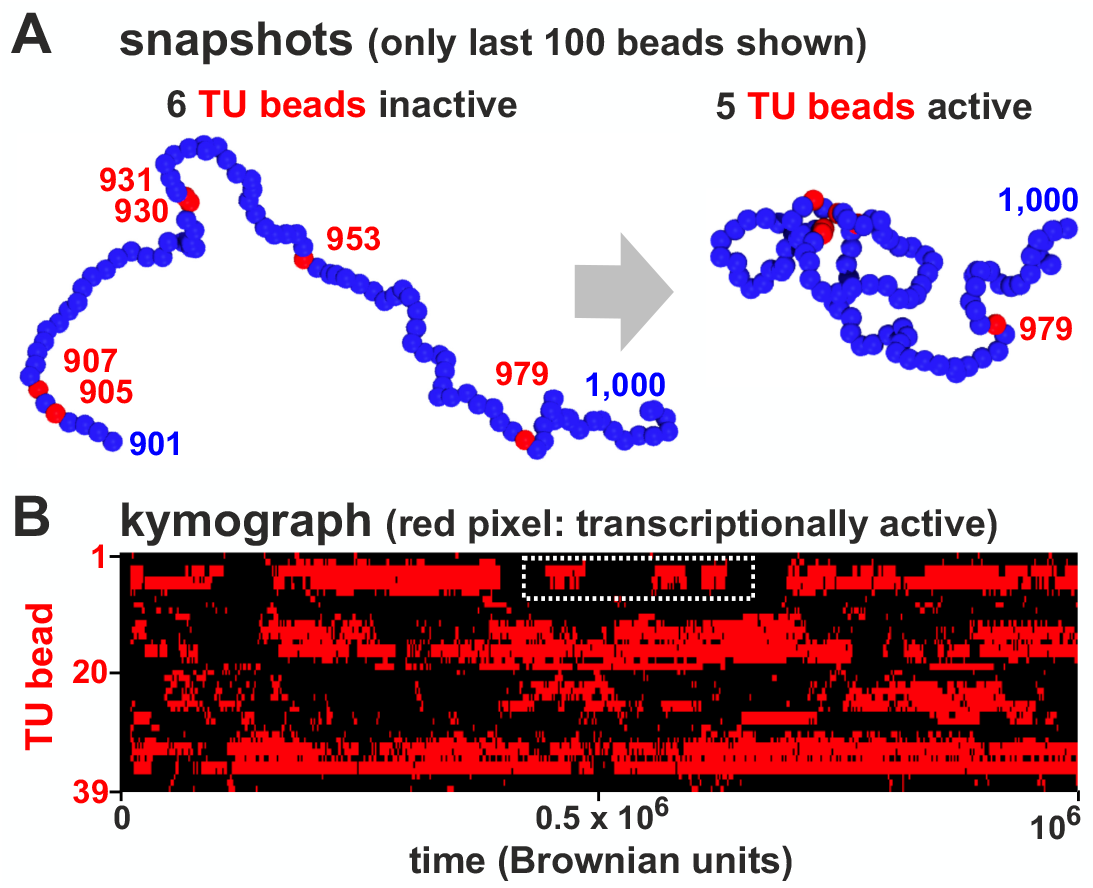
Transcriptional bursting.. **A.** Snapshots showing a 100-bead section of the simulated chain taken at different times. Initially, none of the 5 TUs (red) are in clusters and are inactive; later, 4 TUs join a cluster and are close to TFs – and so are transcribed. **B.** Kymograph where each row shows the changing transcription state of one TU during a simulation; pixels are colored red if the bead is associated with a TF and so transcribed, or black otherwise.

### Local chromatin architecture creates small-world percolating transcription networks

To investigate correlations between transcriptional activities of different TUs, we compute the Pearson correlation matrix between the activity of all possible TU pairs, and identify an emergent regulatory network in which TUs form nodes (Fig. 3A and Fig. S2). Specifically, we draw an edge between two TUs whenever there is a statistically significant positive or negative correlation between their transcriptional dynamics (Fig. 3A). This network arises only due to spatial interactions, as we assume no underlying biochemical regulation.

**FIG. 3:**
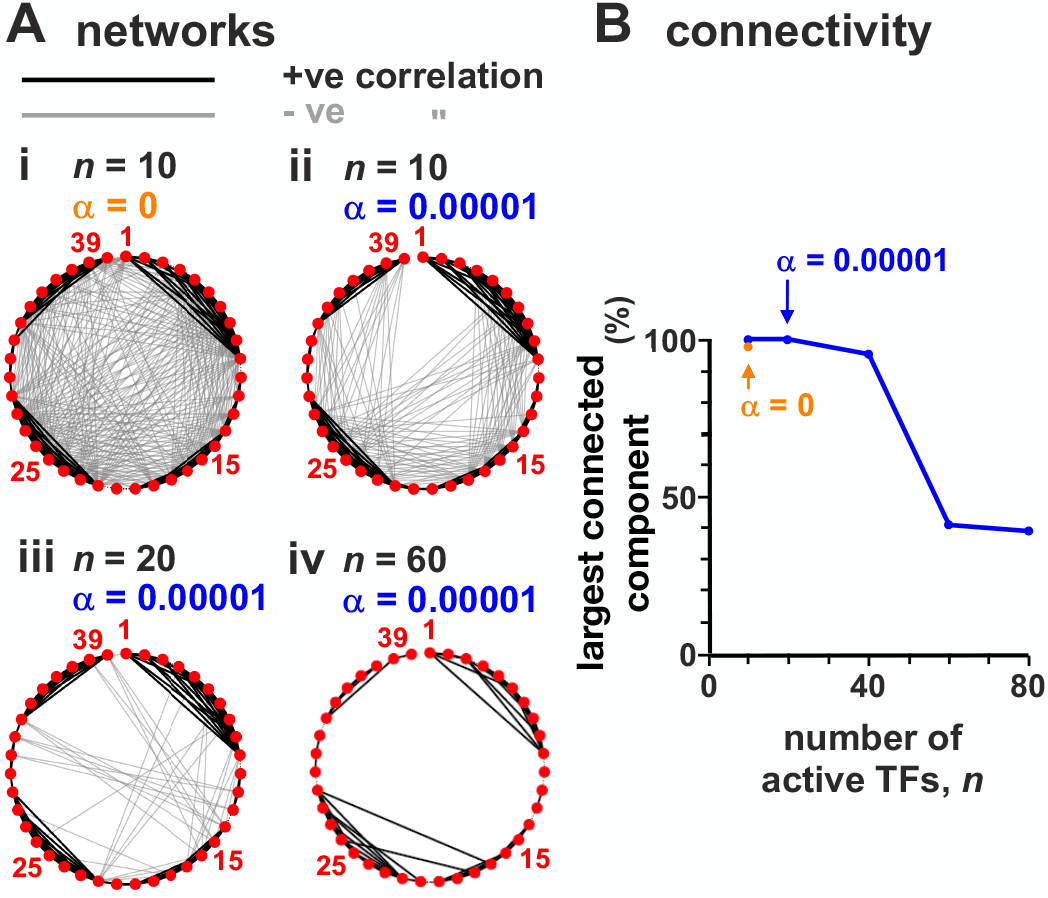
Regulatory networks formed by TU beads are percolating at low TF concentrations. Simulations (as Fig. 1, with ≥ 800 simulations/condition) with different average numbers of active TFs (*n*) and switching rate (*α*). Networks were constructed by calculating the Pearson correlation between the transcription time-series for all pairs of TUs; nodes represent each of the 39 TUs and edges are placed between nodes where there is a significant correlation (> 0.15 in absolute value, corresponding to *p* < 10^−6^). **A.** Effect of TF concentration and switching. 39 nodes are shown around the perimeter, and thick black and grey lines denote positive and negative correlations between transcriptional activities of bead pairs. **B.** Effect of *n* on the (percolating) fraction of nodes in the largest connected component.

The network shows a striking property. With *n* = 10 active TFs, most nodes are connected (Fig. 3Aii), and the fraction of TUs participating in the largest connected component is close to 1 (Fig. 3B). Such a network is said to be “percolating” (any two nodes are connected by a path along edges) and “small-world” (the number of steps between nodes grows as the logarithm of the number of nodes). This phenomenology is consistent with the multitude of small-effect eQTLs detected by GWAS [5, 6]. Notably, these effects act at the transcriptional level, and not post-transcriptionally as envisaged by the omnigenic model [5, 6].

It is important to consider how an apparently simple model can give rise to a complex regulatory network. By analysing simulation trajectories, we noted that TUs lying near each other in 1D sequence space often joined the same cluster in 3D. As a result, the activity of these beads is highly positively correlated (i.e., TUs in the same cluster tend to be active at the same time). At the same time, cluster formation sequesters TFs and so reduces the likelihood that another cluster forms elsewhere. As a result, most long-range correlations are negative (Fig. 3A).

Crucially, these network properties depend on there being a low TF copy-number (as *in vivo* [19]) so that TU beads do not become saturated. We therefore reasoned that increasing copy number should suppress correlations as more rarely-transcribed TUs are pressed into use. Indeed, increasing *n* reduces long-range negative correlations (Fig. 3Aiii,iv), and the fraction of nodes in the largest-connected component falls (Fig. 3B). Another way to think about this result is: if resources are plentiful, there is no need for sharing or competition, and all TUs can bind a TF independently of each other. If TFs do not switch and are permanently in the binding state (and *n* = 10), the network becomes even more highly connected (Fig. 3Ai).

### Modelling eQTL action

GWAS reveals that genetic polymorphisms within eQTLs can lead to many small changes in transcriptional activity across the genome. To model eQTL action, we abrogated TF binding to one TU in the chain; bead 930 was chosen first because it is usually highly active (Fig. 1C). This single “knock-out” affects in a statistically significant way the activity of almost half of the other TUs, both near and far away in sequence space (Fig. 4Aii). The immediately adjacent TU (i.e., bead 931) is down-regulated the most, while more distant ones are up-regulated (due to loss of a strong competitor). This knock-out also rewires the whole network, even though it still retains its small-world character (Fig. 4Aiii). Both positive and negative interactions are affected along the whole chain, as shown by a heat map of the change in Pearson correlation between TU transcriptional activities (Fig. 4Aiv).

**FIG. 4:**
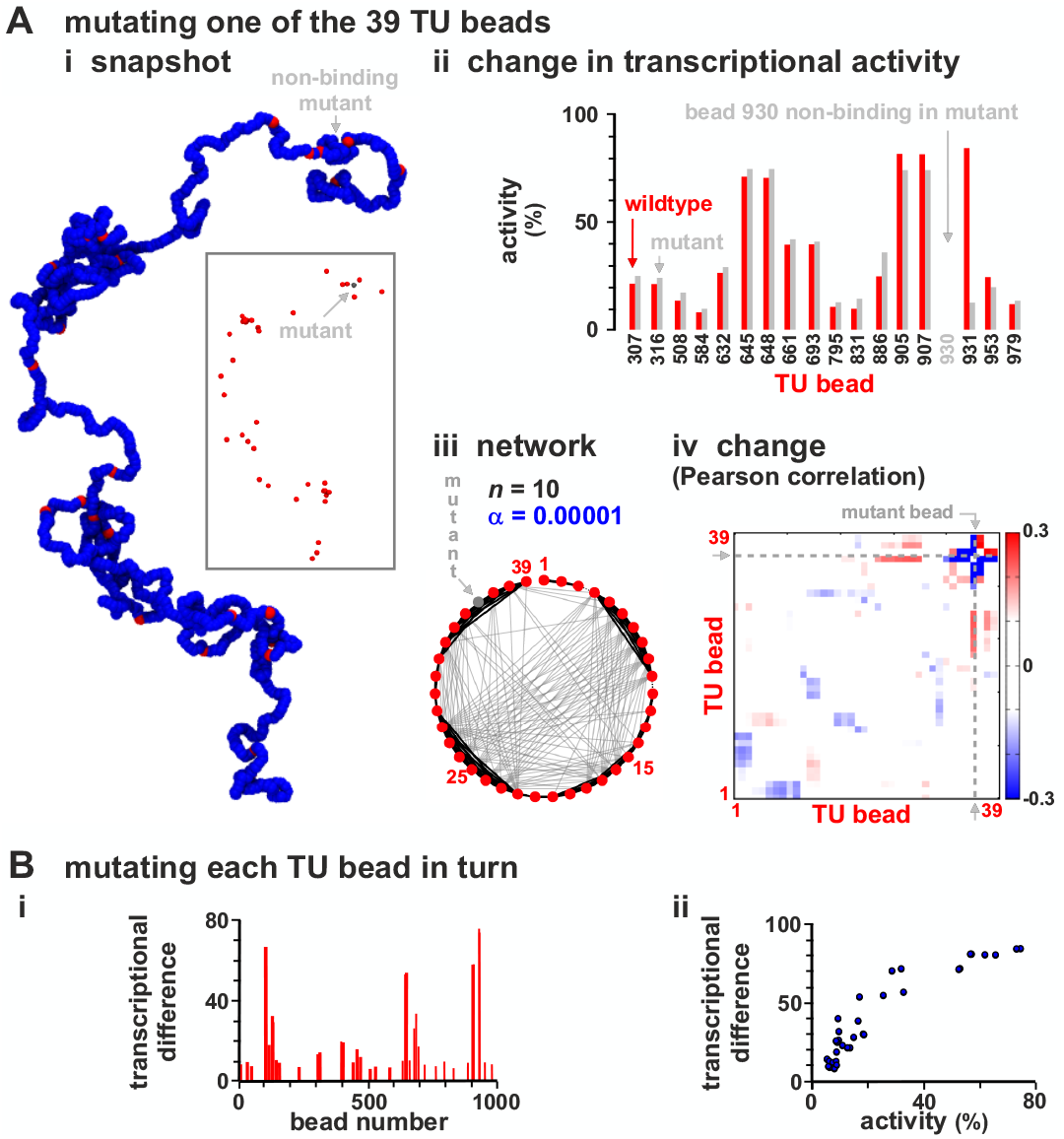
Modelling eQTL action. Sets of simulations (≥ 800 simulations/condition) where each of the 39 TU beads was made non-binding in turn (to represent 39 different eQTLs) are compared with those with the “wild-type” chain (as Fig. 1). **A.** Chain with mutant (non-binding) TU bead 930. (i) Snapshot. TFs not shown (inset: same structure without blue beads). (ii) Transcriptional rates of the 17 TUs with significantly different values in mutant fibre compared with the wild type one (*p* ≃ 0.046; Students t-test). (iii) Regulatory network inferred from the matrix of Pearson correlations between activities of TUs. (iv) Change in Pearson correlation between TUs. **B.** Results from simulations where each TU bead was mutated in turn, and the “transcriptional difference” from the wild type (see text) was determined. (i) Transcriptional difference versus position along the chain. (ii) Positive correlation of transcriptional difference with TU activity in wild-type. The plot therefore shows that if we mutate a TU with high transcriptional activity, this leads to a larger difference.

We next systematically knocked out each TU in turn, one at a time. To quantify global effects, we define a “transcriptional difference” between the wild-type and each knock-out based on a standard Euclidian-distance metric (Eq.(8), Methods); the larger this quantity, the more different the two states are. This difference varies > 10-fold between different mutations (Fig. 4Bi).

Together these observations are reminiscent of the behaviour of eQTLs. In our model, each TU may be seen as an eQTL with different strength, with large-effect eQTLs corresponding to active TUs often in clusters and small-effect ones to TUs which are more isolated in sequence space (Fig. 4Bii).

### Modelling loops, heterochromatin and euchromatin

In mammalian genomes, promoter-enhancer pairs are often contained in loops stabilized by cohesin and the CCCTC-binding factor (CTCF) [31–33]. To investigate how such loops might affect transcription, we incorporated eight permanent and non-overlapping loops at different positions in the chain (Fig. 5A, loops *a-h*). This has subtle but complicated effects. For example, loop *h* encompasses three TUs (beads 905, 907, 930), and expression of one is slightly boosted compared to the unlooped case (Figs. 5B,C). This is consistent with the idea that looping can switch on some genes during development [34]. However, loop *d* encompasses two TUs (beads 396, 404), and has no effect on their activity. Broadly speaking, looping up-regulates activity, but not invariably so, and – perhaps surprisingly – two of the three most up-regulated TUs (beads 33, 886) are not contained in loops (Fig. 5C).

**FIG. 5:**
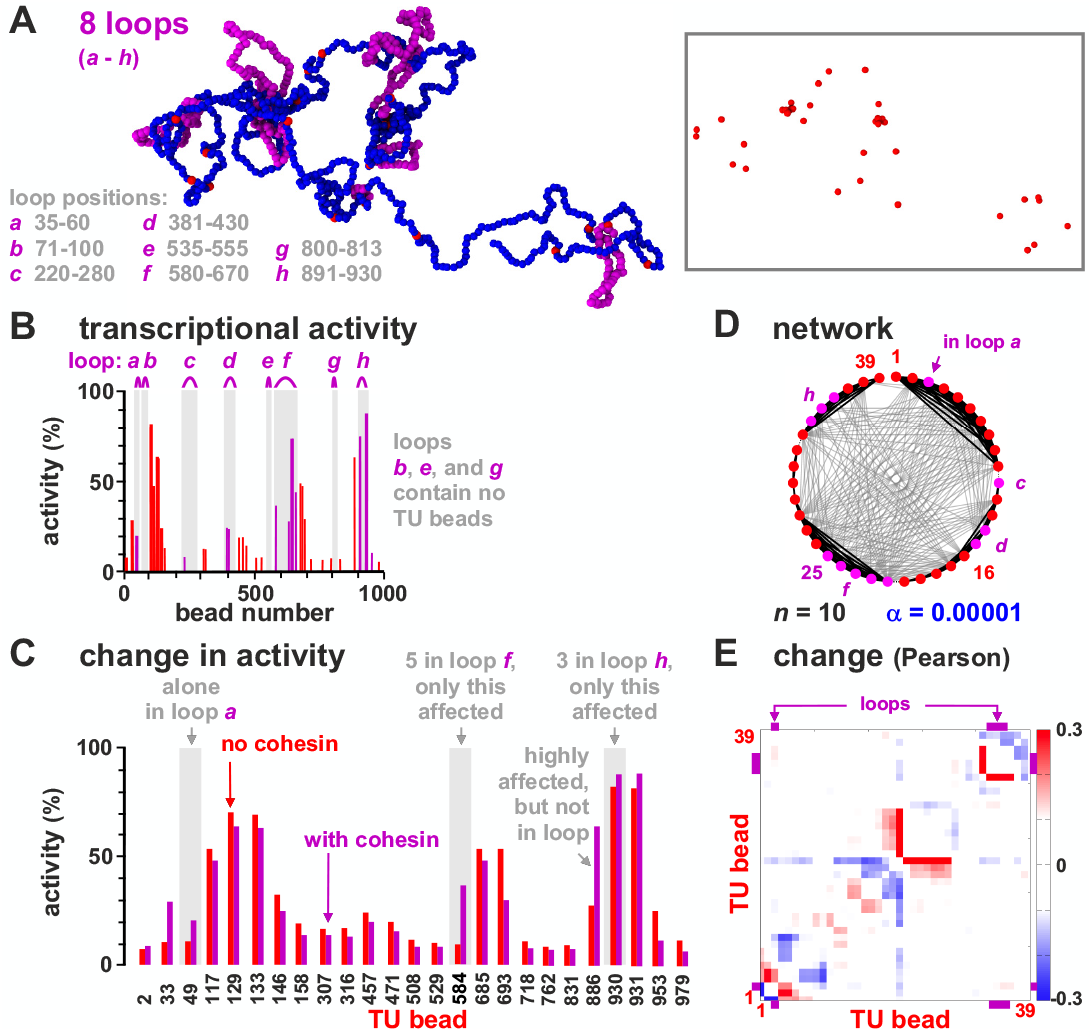
Looping subtly affects transcriptional activity. Results of two sets of simulations were compared; one set was as Fig. 1, in the other the chain contained 8 permanent loops (to represent convergent loops stabilized by cohesin/CTCF). **A.** Snapshot (beads within loops are magenta; TFs not shown; inset – same structure with only TUs shown). **B.** Average transcriptional activity for each TU in the looped chain (magenta bars – values for TUs in loops; magenta arcs – loop positions). **C.** Comparison between activity in wild-type and looped configuration for the 25 TUs with significantly different values in the two sets (*p* ≃ 0.003; Students t-test). **D.** Regulatory network inferred from the matrix of Pearson correlations between expression of TUs (as Fig. 3A). **E.** Change in Pearson correlation between TUs due to loops.

Looping also extensively rewires the regulatory network (Fig. 5D,E). Globally, the increase in activity is modest, as incorporating all beads into closely-packed loops only increases total activity by ~ 10%, with – once again – some TUs being down- as well as up-regulated (Fig. S3). This is consistent with experiments showing that the interplay between looping and expression is complex [35] but slight (e.g., knocking down human cohesin leaves expression of 87% genes unaffected, with global levels changing < 30% [36]).

In all simulations thus far, TFs bind strongly to TU beads, and weakly to all others to model binding to open euchromatin [15, 37]. To investigate the effects of heterochromatin to which TFs bind poorly, and which carries few histone marks [38], is gene poor and traditionally viewed as transcriptionally inert, we performed simulations where four of the most-active TUs (905, 907, 930 and 931) were embedded in a non-binding segment (running from bead 901 − 940). This has a dramatic effect (Figs. 6A-C): the activity of the two TU beads now embedded in the non-binding island are halved, some nearby neighbors are down-regulated, and more distant ones are up-regulated (again due to a reduction in competition; Figs. 6B,C). The regulatory network is also rewired (Figs. 5D,E).

**FIG. 6:**
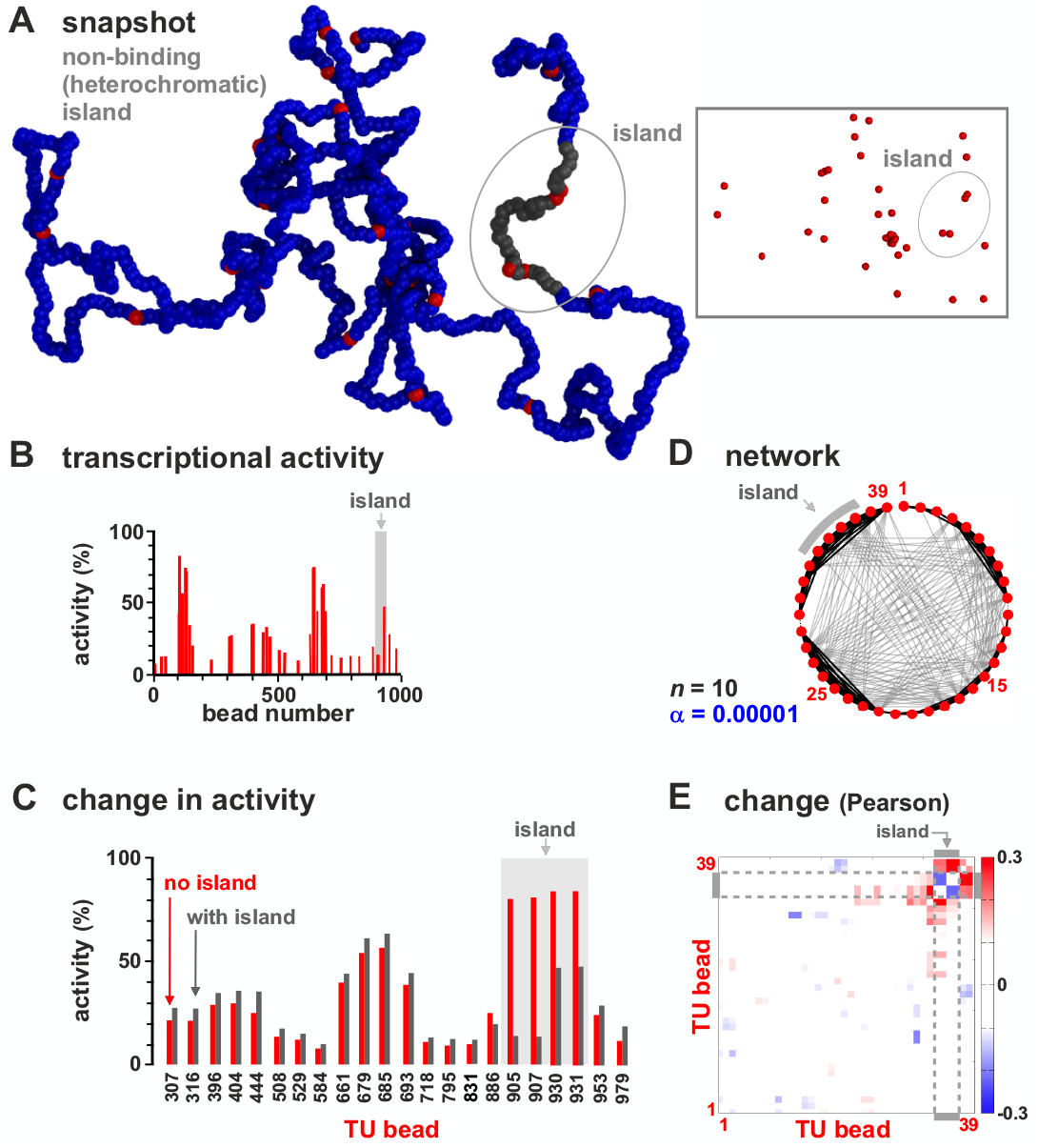
Neighboring heterochromatin affects transcriptional activity. Results from two sets of simulations (at least 800 runs for each condition) were compared; one set was as Fig. 1, in the other beads around TU beads 905, 907, 930 and 931 (from bead 901 to 940) were non-binding (to represent embedding the TU beads in heterochromatin). **A.** Snapshot with heterochromatic beads shown in gray (TFs not shown; inset – the same structure with only TUs). **B.** Average transcriptional activity for each TU. **C.** Comparison of average transcriptional activity with respect to wild-type for the 22 TUs with significantly different values in the two sets (*p* ≃ 0.003; Students t-test). **D.** Regulatory network inferred from the matrix of Pearson correlations between activities of TUs (as Fig. 3A). **E.** Change in Pearson correlation between TUs due to heterochromatin.

Just as an active TU bead can be down-regulated by embedment in a non-binding segment, an inactive one can be up-regulated by embedment in a weak-binding (euchromatic) segment (Fig. S4). This shows our model effectively captures position effects where the local chromatin context strongly influences activity [39].

### Modelling a whole human chromosome

We next modelled a whole mid-sized human chromosome (HSA 14, length 107 Mbp; Fig. 7A), in a well-characterized and differentiated diploid cell (HUVEC, human umbilical vein endothelial cell). Now, each bead (again with diameter of 30 nm) in a chain of 35784 beads represents 3 kbp in the chromosome, and we used experimental data to identify TUs as well as euchromatic and heterochromatic regions. As before, multivalent TFs are present at a non-saturating concentration and switch between active and inactive states (with 20% active at any time). TFs bound strongly to TU beads, weakly (non-specifically) to euchromatin beads, and had no affinity for heterochromatin beads. As chromosome territories are often ellipsoidal, simulations are performed in an ellipsoid of appropriate size [7, 40]; consequently, chromatin density is higher than in the simulations detailed above, with volume fractions comparable to those *in vivo* (~ 14%).

**FIG. 7:**
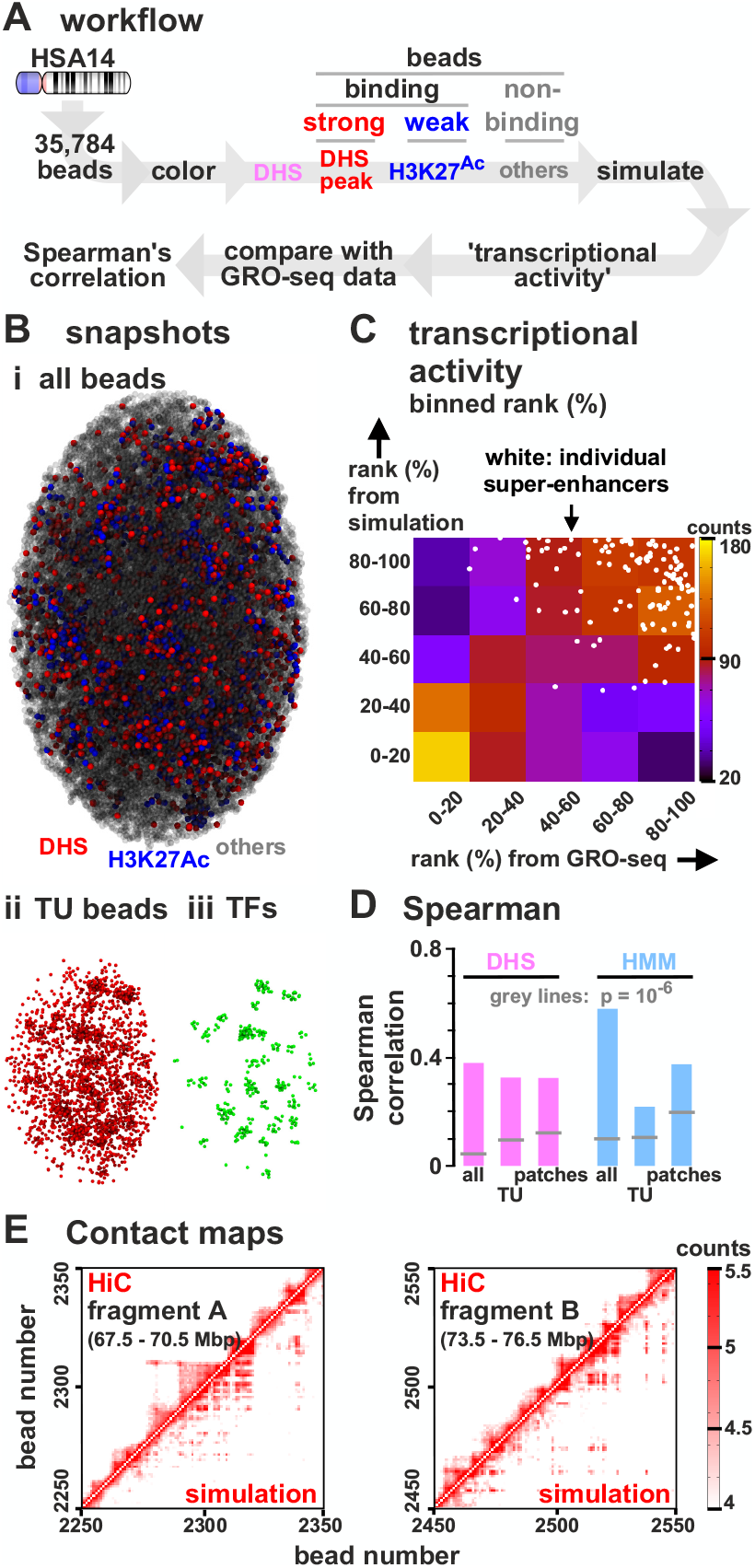
Comparison of transcriptional activities of TUs on HSA14 in HUVECs determined using simulations and GRO-seq. **A.** Workflow (DHS model). Simulations (244 runs) involved a chain (35784 beads) representing HSA14, and 1700 switchable TFs confined in an ellipsoidal territory. Beads were classified as TUs (red, strong-binding), euchromatic (blue, weak-binding) or heterochromatic (grey, non-binding). Transcriptional activities from simulations are compared with those of GRO-seq data, by measuring the Spearman rank correlation. **B.**(i) Snapshot (TFs not shown). (ii, iii) TU beads and TFs in this configuration. **C.** Comparison of transcriptional activities of TUs from simulations and GRO-seq (ranked from 0 − 100%, then binned in quintiles). A scatter plot of unbinned ranks of beads corresponding to SEs are superimposed (white circles). **D.** Comparison of transcriptional activities from simulations (for both DHS and HMM models) and GRO-seq for all 3 kb regions/beads, only TUs, and only connected patches of binding beads (see text). All correlations are significant (*p* < 10^−6^, indicated by grey lines). **E**(i-ii) Capture-HiC-like contact maps obtained from simulations and experiments [31] showing contacts between 30 kbp bins which contain TUs.

The types of chromatin beads were identified using DNase-hypersensitity data and ChIP-seq data for H3K27ac. DNase-hypersensitive sites (DHS) are excellent markers to locate promoters and enhancers, and in general TF binding sites [15, 41], whereas H3K27ac modifications strongly correlate with open chromatin [15]. Therefore, if the 3 kbp region corresponding to a chromatin bead has a DHS, then that bead is a TU; if it has H3K27ac, it is a euchromatin bead – and all other beads are non-binding (and so heterochromatic). We call this the “DHS” model. As properties of different chromatin segments have been catalogued using “hidden-Markov models” (HMMs) applied to many data sets [38], we can alternatively classify beads according to HMM state of the corresponding chromosomal region; we call this the “HMM model”.

Simulations using the DHS model again yield clusters enriched in TUs and TFs (Fig. 7B). As before, aggregating data from many simulations allows determination of transcriptional activities of every bead, which we compare with those of corresponding regions determined experimentally [42] by GRO-seq (global run-on sequencing [43]); activities of all 3 kbp regions were ranked from high to low, binned into quintiles, and compared. In Figure 7C, squares near the diagonal from bottom-left to top-right have high ranks (shown as red and yellow) compared to those off-diagonal (blue and purple) indicating good concordance between simulations and data. A specific sub-set of beads corresponding to super-enhancers (SEs) – which are highly active *in vivo* [44] – are also highly active in simulations (shown as white dots concentrated at top right). This concordance between results from simulations and GRO-seq is confirmed by the Spearman rank correlation (~ 0.38 for all beads; *p* < 10^−6^; this measure is used because it is less sensitive to outliers; Fig. 7D). Restricting analysis just to TUs provides a more stringent comparison (as all TUs bind TFs with equal affinity); it still yields a significant correlation (*r* ≃ 0.32, *p* < 10^−6^; Fig. 7D). As neighbouring high-affinity regions tend to have roughly similar transcriptional rates in both simulations and data, we also average rates found in active “patches” (contiguous sets of beads which are either all TUs or all labelled as euchromatin), but found this has no significant effect (Fig. 7D). Concordance was confirmed using our HMM model (Fig. 7D, right, and Fig. S5). Adding cohesin-mediated looping to simulations involving the DHS model did not significantly change agreement with experimental data (e.g., for TUs only, *r* ≃ 0.33, *p* < 10^−6^). Similar agreement with GRO-seq data was obtained from simulations applied to the H1 human embryonic stem-cell line (for TUs using the DHS model, *r* ≃ 0.29, *p* < 10^−6^).

Again, regulatory networks are small-world and highly connected (Fig. S6). To facilitate comparison with previous results, four segments of HSA14 were selected that had the same length as the toy one (i.e., 3 Mbp), and roughly the same density of TUs; all four segments again had highly-connected components (compare Figs. S6 and Fig. 4). However, patterns in real chromosomes and artificial fragments are quite different. In HSA14 networks, there are more positive interactions between sets of adjacent TUs and other sets that are > 10 beads distant in sequence space (black lines across the middle of circles in Fig. S6).

We previously showed [12] that simulations involving two different TFs (that bind to active and inactive regions, respectively) yield contact maps much like those found with Hi-C [31]. Therefore, we expected the present simulations to reflect Hi-C data poorly as they involve only one TF binding to the minor (i.e., active) fraction of the genome, so contacts made by this structured minority would be obscured by those due to the unstructured majority. Even so, simulations yield contact maps broadly similar to those obtained by Hi-C with excellent Pearson correlations (*r* = 0.93, *p* < 10^6^; not shown). To focus on the structured minority, we use a more relevant comparison based on contact maps restricted to TUs as anchors – which may be considered as equivalent to interactions obtained by promoter-capture HiC [45]. These yield good concordance (Fig. 7E; Pearson coefficient *r* = 0.82, and *r* = 0.47 when monitoring only long-range contacts between TUs at least 300 kbp away; *p* < 10^−6^).

Overall these results show that a simple model based on 3D chromatin organisation can capture much of the complexity in transcriptional dynamics of a whole human chromosome.

### Modelling chromosome 22 carrying the diGeorge deletion

Our approach can, in principle, be applied to study any chromosome providing appropriate genomic data are available (e.g., on DNase hypersensitivity and histone acetylation). As a proof of principle, we studied the effect of deleting ~ 2.55 Mbp from HSA22 – an alteration which is associated with the diGeorge syndrome (Fig. 8A) [52]. This syndrome affects ~ 1 in 4000 people, and the variable symptoms include congenital heart problems, frequent infections, developmental delays, and learning problems. We predict a multitude of small effects in TU activity, both near and far away from the deletion (see the Manhattan plot in Fig. 8Bi). In particular, most TUs are slightly up-regulated, as fewer TUs compete for the same number of factors, and the TUs which change the most are those corresponding to intermediate transcriptional activities in the wild-type (Fig. S7). The p-values associated with the change in transcriptional activities vary widely, and comparison of the observed distribution with the null hypothesis (indicating that changes in measured transcription are due to random variation) shows the observed is highly enriched in small p-values (Fig. 8Bii), as is generally the case with results from GWAS [5, 6]. The regulatory network is also re-wired (Fig. 8C).

**FIG. 8:**
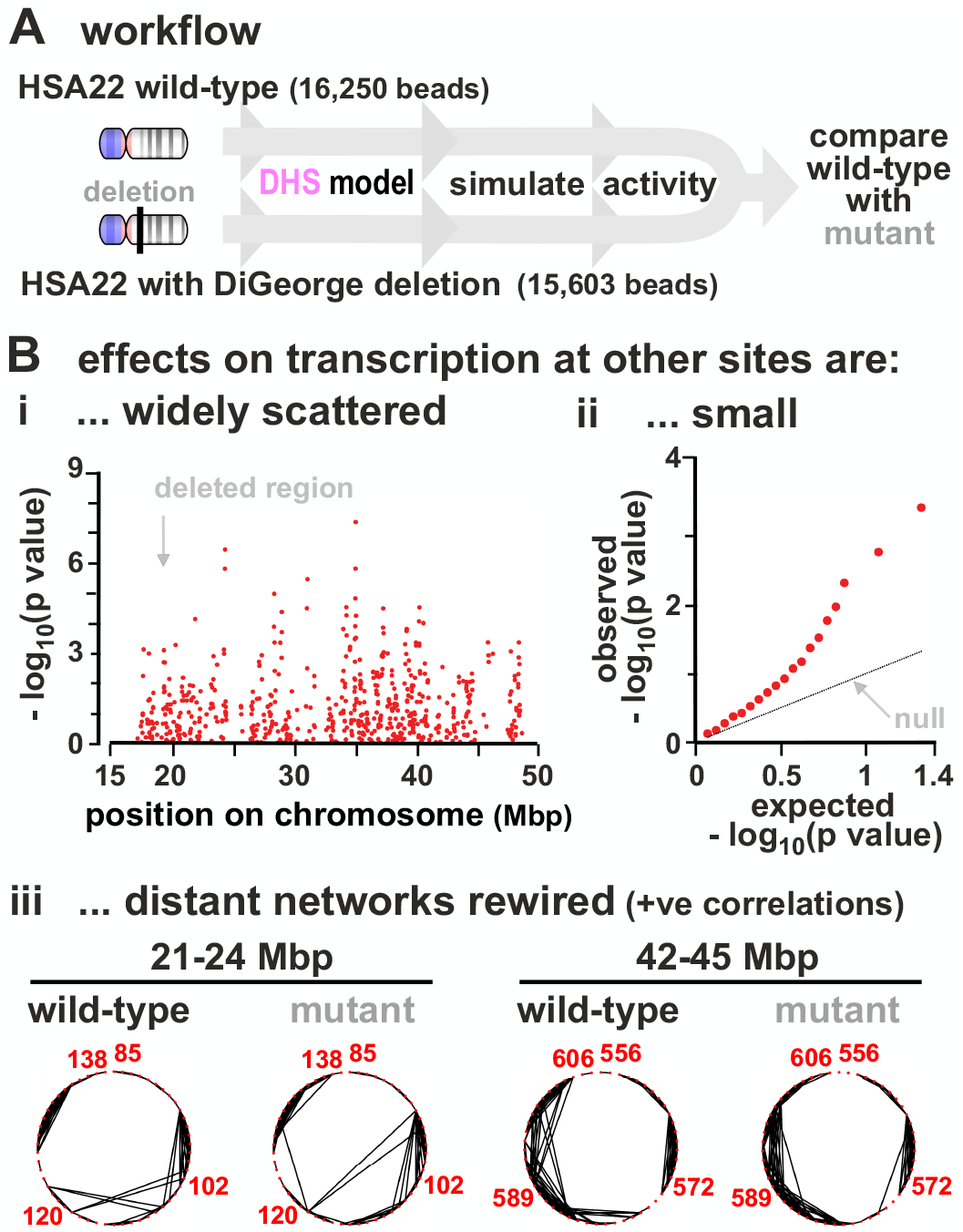
Modelling effects of the DiGeorge deletion in HSA22. **A.** Workflow. Simulations (800 simulations/condition) for the wild type (17102 beads) and deletion (16250 beads, where wild-type beads 6305 − 7156 were cut, corresponds to a deletion of chr22:18912231 − 21465672 in hg19). **B.** (i) Manhattan plot showing log_10_(p-value) as a function of genomic position along HSA22 (position given in Mbp), for changes in TU transcriptional activities between wild-type and deletion. (ii) Quantile-quantile plot showing expected versus observed values for log_10_(p-value) for the same data in (i). Expected values were computed from the normal distribution (these correspond to the null hypothesis according to which the change in transcriptional activities in the deletion is purely due to random variation). **C.** Regulatory networks of two 3 Mbp segments in chromosome 22 inferred from the Pearson correlation matrix. Edges show positive correlations > 0.12 (*p* = 0.0007). Segments were chosen because they have roughly the same number of nodes in 3 Mbp as the short fragment (Fig. 3Aii).

Clearly, this approach opens up a rich new field of study. For istance, while there may be processes which occur *in vivo* which are not represented in the current model, it could still give an indication of the genes most likely to be affected by any chromosome rearrangement.

## DISCUSSION AND CONCLUSIONS

We have described a parsimonious 3D stochastic model for transcriptional dynamics based just on multivalent binding of factors and polymerases (TFs) to genic and nongenic transcriptional units (TUs) in a chain representing a chromatin fibre. Two different fibres were considered: a 3 Mbp fragment with randomly-positioned TUs, and human chromosome 14 where TUs were appropriately positioned along 107 Mbp. Despite deliberately excluding any explicit underlying network of biochemical regulation, our model nevertheless yields several striking results. These depend on having a low TF copy-number – a feature compatible with observations *in vivo* [19]. First, since TFs bind with the same affinity to all TUs, one might expect all TUs to in 1D sequence space tend to be the most active (Fig. 1C) with positively-correlated dynamics reminiscent of transcriptional bursting (Fig. 2B). This is because they often cluster into structures analogous to the phase-separated transcription hubs/factories seen experimentally [7, 10]. Second, switching off binding at any TU significantly affects the activity of many others, both near and far away in sequence space (Fig. 4). Third, introducing stable loops into the fibre has only modest effects on activity (Fig. 5), consistent with the modest reductions seen experimentally in cohesin knock-outs [36]. Fourth, transcriptional activity of a TU is strongly affected by the local environment in ways that are reminiscent of the silencing of a gene by incorporation into heterochromatin [39] (Fig. 6), or activation by embedment in euchromatin (Fig. S4). Finally, our simple fitting-free model captures much of the rich complexity seen in the transcription of a whole human chromosome (Fig. 7).

Our most important result is that our simulations reveal small-world networks of mutual up- or down-regulation between TUs (Figs. 3 and S6). These networks provide a framework in which to understand how regulation of any particular gene is determined by a multitude of eQTLs, each having only a tiny effect [5, 6] – a result which is difficult to explain with any model that disregards spatial effects (as each interaction would then require some specific biochemical regulation). At the same time, we show that increasing TF copy-number provides a simple but effective way to dramatically simplify network structure by removing correlations (Fig. 3). We suggest this occurs when a fibroblast is reprogrammed into a muscle cell by flooding cells with the MYOD factor, or into pluripotent stem cells by the four Yamanaka factors. In other words, overexpressing TFs over-rides the complex regulatory network mediated by 3D structure, and allows the coexisting transacting biochemical network to dominate.

Taken together, these results suggest the activity – or inactivity – of every genomic region affects that of every other region to some extent. We describe our framework as “pan-genomic” (Fig. S8). This is reminiscent of the omnigenic model [5, 6] in the sense that many loci are involved, all having small effects. However, our model differs in several major respects. It incorporates genome organisation and gene regulation and gives molecular detail on both (the omnigenic model says little about organisation, and is openly short on molecular detail [6]). It is more extreme as all genomic regions contribute; non-genic transcription units play even more important roles than genic ones as they are more numerous, with chromosome loops, heterochromatin, and euchromatin all playing additional roles. Moreover, it posits a direct and immediate effect of structure on regulation at the transcriptional level, which contrasts with the non-trivial post-transcriptional pathways envisioned by the omnigenic model.

Our framework could be applied to predict the transcriptional activity of any genomic fragment, or the effect of any chromosome deletion (Fig. 8) or rearrangement. Its predictive power could be improved further by incorporating data on histone modifications in the wild-type and mutant genomes. For instance, it could be employed to predict the transcriptional changes arising from 3D structure following rearrangements occurring in cancer.

We thank the European Research Council (ERC CoG 648050 THREEDCELLPHYSICS) for support.

## METHODS

### Polymer modelling

In this work we model chromatin fibres and chromosomes as bead-and-spring polymers. A fibre has *M* monomers, each of size *σ* (corresponding to 3 kbp, or 30 nm [20]), and **r**_*i*_ denotes the position of the *i*-th monomer in 3D space. Multivalent transcription factors (either active or inactive) are also modelled as spheres, again with size *σ* for simplicity. There are *n* multivalent factors in a simulation (where *n* was varied systematically, see text and Results section for details), and *N* high-affinity binding sites, which we refer to as transcriptional units, or TU (or TU beads).

Any two monomers (*i* and *j*) in the chromatin fibre interact purely repulsively, via a Weeks-Chandler-Anderson potential, given by

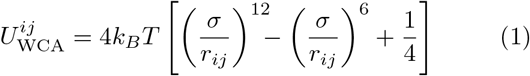

if *r_ij_* < 2^1/6^*σ* and 0 otherwise, where *r_ij_* is the separation of beads *i* and *j*. There is also a finite extensible non-linear elastic (FENE) spring acting between consecutive beads in the chain to enforce chain connectivity. This is given by

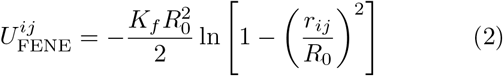

where *i* and *j* are neighbouring beads, *R*_0_ = 1.6*σ* is the maximum separation between the beads, and *K_f_* = 30*k_B_T/σ*^2^ is the spring constant.

With simulations including permanent cohesin loops (Fig. 7 and Suppl. Fig. S4), neighbouring monomers and monomers forming loops interact via harmonic, rather than FENE springs,

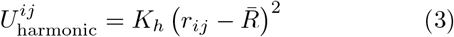

where *i* and *j* are neighbouring beads, *K_h_* = 100*k_B_T/σ*^2^ is the harmonic spring constant, and 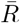 is the equilibrium spring distance. For these simulations, we used 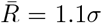 for bonds joining neighbouring monomers along the chain, and 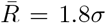 for bonds joining loop-forming monomers. The harmonic potential was used instead of the FENE one to enhance numerical stability.

Finally, a triplet of neighbouring beads interact via a Kartky-Porod term to model the stiffness of the chromatin fibre. This term explicitly reads as follows,

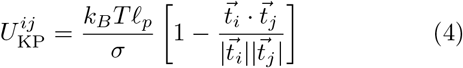

where *i* and *j* are neighbouring beads, 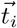 is the tangent vector connecting beads *i* to *i* + 1, and *l_p_* is related to the persistent length of the chain: this parameter is set to 3*σ* in our simulation, which corresponds to a relatively flexible fibre – the resulting persistence length is within the range of values estimated for chromatin from experiments and computer simulations [46].

The interaction between a chromatin bead, *a*, and a multivalent TF, *b*, is modeled through a truncated and shifted Lennard-Jones potential, given by

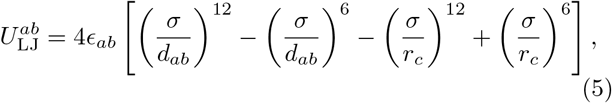

for *d_ab_* (the distance between the centres of chromatin bead and protein) smaller than *r_c_*, and 0 otherwise. The parameter *r_c_* is the interaction cutoff; it is set to *r_c_* = 2^1/6^*σ* for inactive proteins or for active proteins and non-binding chromatin beads (this cutoff results in a Weeks-Chandler-Anderson potential and purely repulsive interactions), or to *r_c_* = 1.8*σ* for an active protein and a sticky chromatin bead (this results in an attractive interaction to model binding). In all cases, the potential is shifted to zero at the cut-off in order to have a smooth potential. Purely repulsive interactions are modeled by setting *E_ab_* = *k_B_T*, while attractive interactions are modeled using *E_ab_* = 3*k_B_T* for active TF and low-affinity beads, and to *E_ab_* = 8*k_B_T* for active TF and high-affinity (TU) beads.

A TU bead (or more generally any chromatin bead in Fig. 8D) is said to be transcribed if it is bound to a factor – i.e., if there is at least a TF whose centre lies within a range *r_c_* = 1.8*σ* away from the bead centre.

The time evolution of each bead in the simulation (whether TF or chromatin bead) is governed by the following Langevin equation

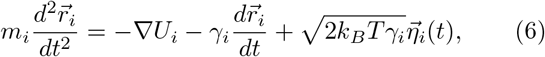

where *U_i_* is the total potential experienced by bead *i*, *m_i_ ≡ m* and *γ_i_ ≡ γ* are its mass and friction coefficient (equal for all beads in our simulations), and 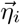 is a stochastic noise vector with the following mean and variance

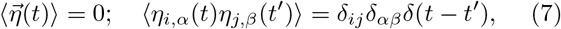

where the Latin and Greek indices run over particles and Cartesian components, respectively, and *δ* denotes here the Kronecker delta.

As is customary [47], we set *m/ξ* = *τ*_LJ_ = *τ*_B_, with the LJ time 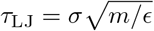 and the Brownian time *τ*_B_ = *σ/D*, where *ϵ* is the simulation energy unit, equal to *k_B_T*, and *D* = *k_B_T/γ* is the diffusion coefficient of a bead of size *σ*. From the Stokes friction coefficient for spherical beads of diameter *σ* we have that *ξ* = 3*πη*_sol_*σ* where *η*_sol_ is the solution viscosity. One can map this to physical units by setting the viscosity to that of the nucleoplasm, which ranges between 10 − 100 cP, and by setting *T* = 300 K and *σ* = 30 nm, as above. This leads to *τ*_LJ_ = *τ*_B_ = 3*πη_sol_σ*^3^/*E* ≃ 0.6 − 6 ms. The Brownian time *τ*_B_ is our unit of time in simulations. The numerical integration of Eq. (6) is performed using a standard velocity-Verlet algorithm with time step Δ*t* = 0.01*τ*_B_ and is implemented in the LAMMPS engine [48]. Protein switching is including by stochastically changing the type of TF beads every 10000 timesteps (equivalently, every 100 Brownian times), with probabilities such that the switching off rate is of 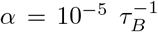, or 0.017 0.17 s^−1^. In simulations of the toy model (Figs. 1-7 and Suppl. Figs. S1-S4), the switching on rate is equal to *α*; in chromosome 14/22 simulations (Fig. 8 and Suppl. Fig. S5), it is equal to *α*/4. Consequently, in steady state the average number of active and inactive proteins is equal in simulations of the toy model, whereas the average number of inactive proteins is 4-fold larger than that of active proteins in chromosome 14/22 simulations.

### Additional simulation details

#### Artificial chromatin fragment simulations

For the 3 Mbp chromatin fragment simulations, chains were made up of 1000 beads (so that each bead corresponds to 3 kbp). For the simulations shown, TUs were placed at beads 2, 33, 49, 103, 105, 117, 129, 133, 146, 158, 233, 307, 316, 396, 404, 444, 457, 471, 508, 529, 584, 632, 645, 648, 661, 679, 685, 693, 718, 762, 795, 831, 886, 905, 907, 930, 931, 953, 979. Different simulations with different TU bead locations (randomly scattered along the fibre with average distance of 30) gave qualitatively similar resuls to the ones shown in the main text.

To start simulations, the chromatin fibre was initialised as a random walk, and proteins were initialised at random position within the simulation box – a cube of size 100*σ*, which means the system is dilute. To avoid bead overlap, we initially perform a small number of steps with a soft potential between beads, and a harmonic bond between neighbouring beads. We then equilibrate the system for 10^4^ Brownian times with repulsive (WCA) interactions between any pair of beads. Finally we study the evolution of the system for 10^5^ Brownian times (10^6^ Brownian times for Fig. 2) once the attractive Lennard-Jones interactions between chromatin beads and TFs are included in the potential.

#### Transcriptional difference

To determine the difference between two sets of simulations in Figure 4, which we refer to as states (typically corresponding to two different parameter sets), we define a transcriptional difference as follows. If there are *N* TU beads in the polymer, let us call *x_i_* the expression (transcriptional activity, in %) of the *i*-th TU bead in the first state, and 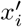 its expression in the second state. The transcriptional difference between the two states is then

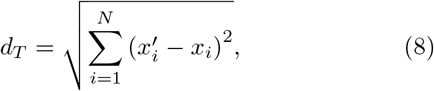

which is the Euclidean distance between the two points {*x*_1_*, · · ·, x_N_*} and 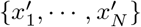 in their *N*-dimensional space.

#### Whole chromosome simulations

To simulate HSA 14 and HSA22 in a diploid G0 HUVEC cell, the system is confined into an ellipsoidal territory with aspect ratio chosen according to typical experimental values [7, 40] (semiaxes were 22.24*σ*:34.24*σ*:41.80*σ*, or 0.67 *μ*m:1.03 *μ*m:1.25 *μ*m, for HSA14; 17.39*σ*:26.77*σ*:32.68*σ*, or 0.52 *μ*m:0.80 *μ*m:0.98 *μ*m, for HSA22). Confinement is enforced by modifying the source code in LAMMPS to describe an ellipsoidal indenter. This introduces a soft force towards the centre of the ellipsoid, only if beads exit the confining ellipsoid. The magnitude of the confining force felt by a bead at position vector (*x, y, z*) is

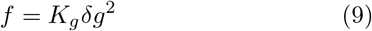

for *δg* > 0, where

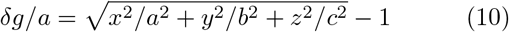

where *a*, *b* and *c* are the semiaxes (along *x*, *y* and *z*), and where we have assumed for simplicity that the centre of the ellipsoid is in (*x, y, z*) = (0, 0, 0). The constant *K_g_* measures the stength of the confining force and was set to 10*k_B_T/σ*^2^ in simulations. As a detail we note that the confinement is modelled via an effective force, rather than via an effective potential. Polymer size (in beads) for the simulated chromosomes were 35784 for HSA14, 17102 for wild-type HSA22, and 16250 for HSA22 with diGeorge deletion (again, each bead corresponds to 3 kbp).

There are 320000 pol II molecules in a nucleoplasmic volume of 660 *μ*m^3^ in HeLa cells [49], equivalent to a 0.8 nM concentration and FRAP shows that about a quarter exchanges rapidly [50] and is likely to be active at any time. To reflect this situation, we considered 1700 and 813 active factors (representing, as usual, complexes of pol II and transcription factors) for HSA14 and HSA22 simulations respectively, with *k*_off_ = 4*k*_on_ = 0.001 *τ*_B_, so that 20% of these are active on average at any time.

### Genomic datasets used

In this Section we detail the genomic datasets used to prepare inputs and compare outputs in simulations of chromosome HSA 14 and 22 (Fig. 8 and Suppl. Fig. S5).

Inputs to color beads in the DHS model are based on DNase-hypersensitivity (DHS) and H3K27ac peaks for HUVECs and hESCs, from ENCODE. Beads containing a peak in H3K27ac but not in DHS are colored as low-affinity (blue) beads, and those containing a peak in DHS are colored as high-affinity (red) beads. Beads which did not contain any peak (either DHS or H3K27ac) are colored as non-binding (gray). To color beads in the HMM model (HUVECs only), we used the chromatin states in [38] as detailed in the main text. Our simulations are based on the GRCh37-hg19 genome assembly for inputs.

To compare the predicted transcriptional activity of chromosome 14 outputted by our simulations with experiments, we use GRO-seq data. For HUVECs, we use the datasets GEO: GSM2486801, GSM2486802, GSM2486803 [42]; for hESCs, we use GEO: GSM1579367, GSM1579368 [51]. Super-enhancer regions considered here are those identified in [44], and available in the dbSUPER database, which includes super-enhancers for human and mouse cells.

---

**FIG. S1:**
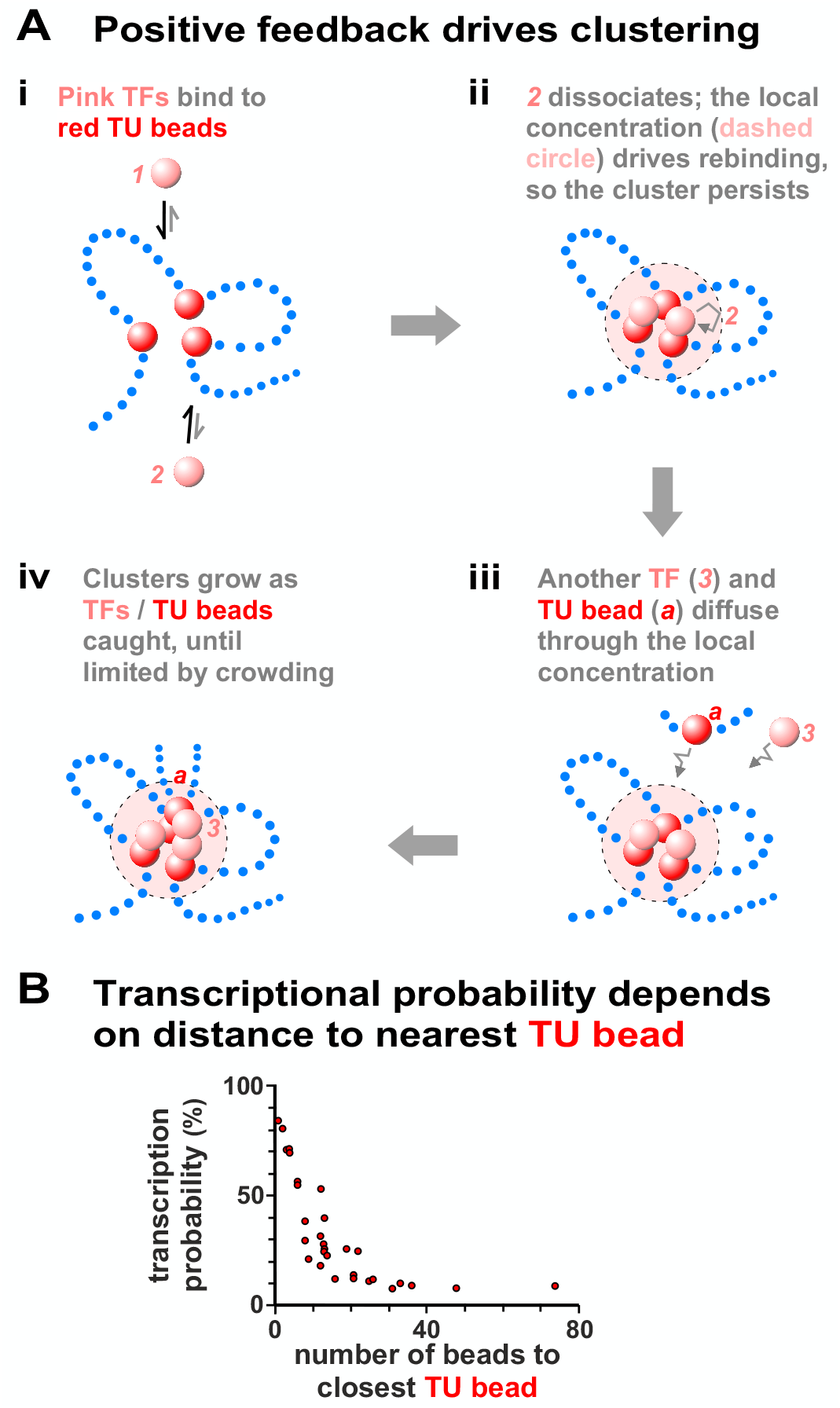
Clusters form spontaneously, and nearby TU beads tend to be transcribed more often. **A.** Clustering is driven by positive feedback (the “bridging-induced attraction”) [12, 20]. (i) Multivalent TFs *1* and *2* (pink) bind reversibly to red TU beads; each blue dot represents many non-binding beads. No other attractive forces between TFs or between TUs are specified. (ii) The two TFs bound; each stabilizes a loop. The local concentration of red TUs in the dashed volume has now increased, so if pink TF *2* dissociates it is likely to rebind to the same cluster (grey arrow); therefore, the cluster is likely to persist. (iii) The high local concentration also drives cluster growth. Here, the cluster will catch red TU *a* and pink TF *3*, as they diffuse through this local region. (iv) Cluster growth continues due to this positive feedback until limited by the entropic costs of crowding together ever-more loops. **B.** Scatter plot where each point represents a TU bead, and the horizontal axis gives the distance along the chain in beads to the nearest neighbouring TU. A strong anticorrelation with transcriptional activity is evident. The positive feedback described in (A) ensures that the closer a TU is to another TU, the higher the probability it will cluster and be transcriptionally active.

**FIG. S2:**
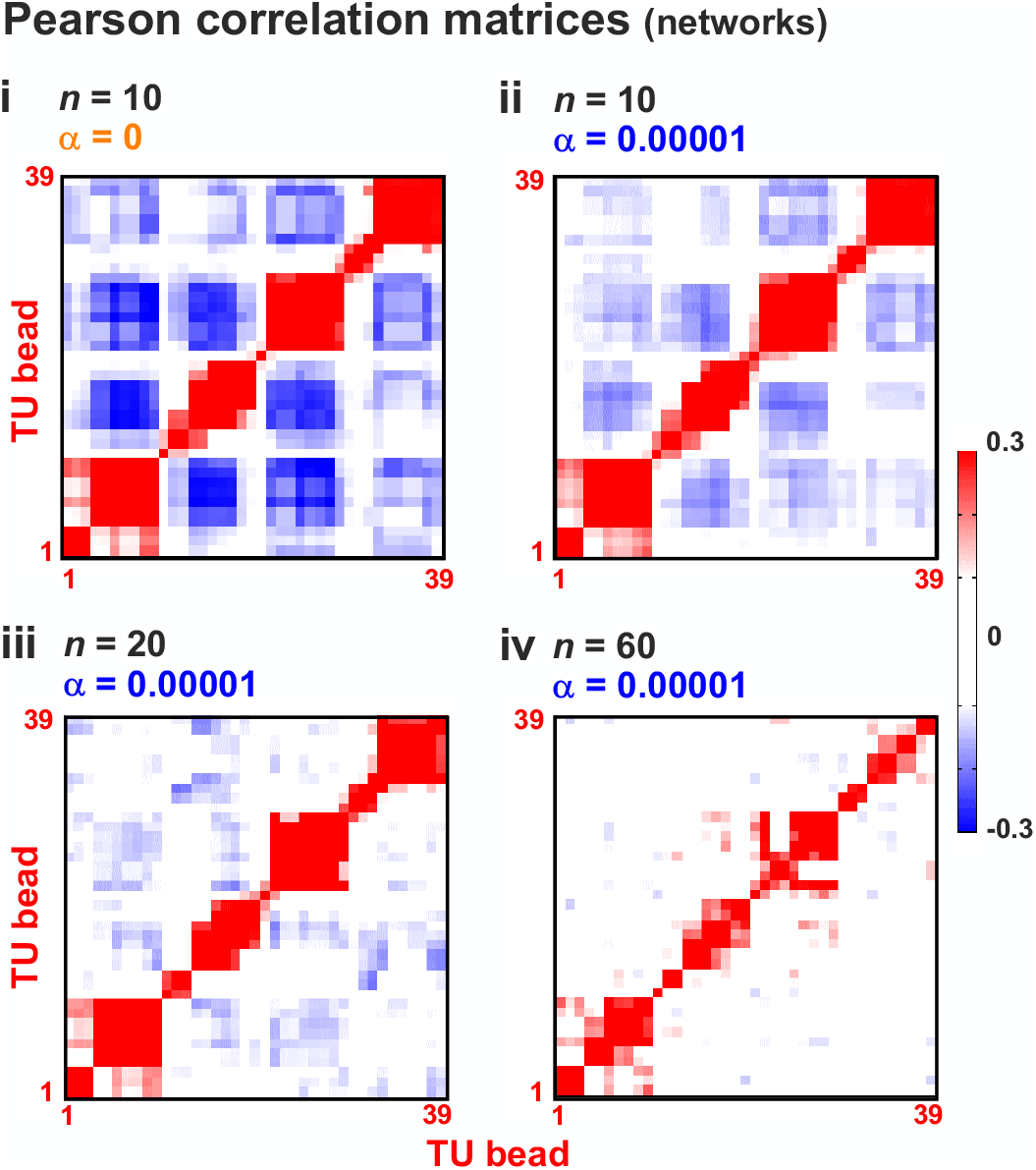
Pearson correlation matrices. Pearson correlation (covariance) matrices for simulations in Fig. 3. These were used to construct regulatory networks.

**FIG. S3:**
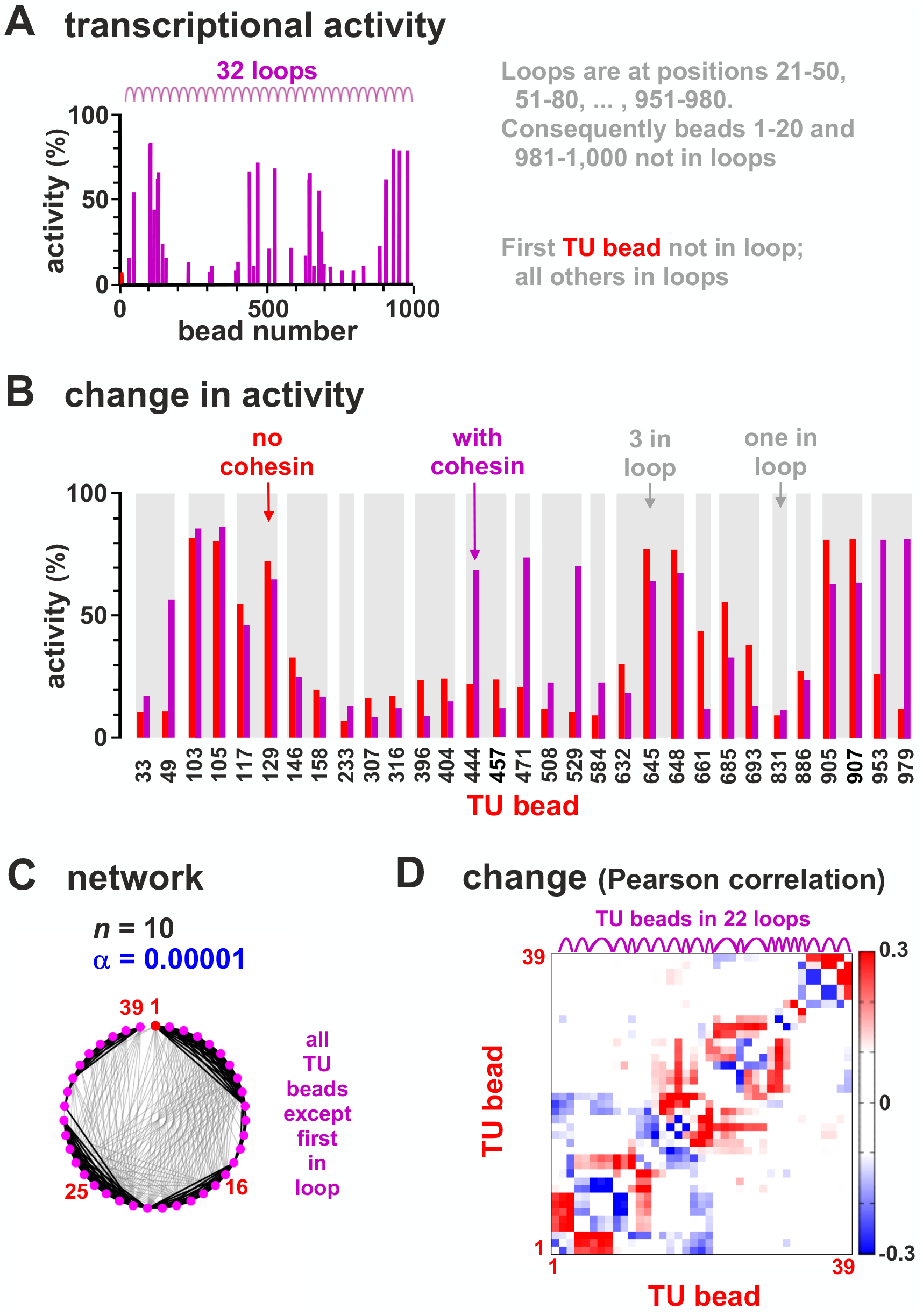
Transcriptional activity in a chain with closely-packed loops. Results of two sets of simulations (800 simulations/condition) were compared; one set was as Fig. 1, in the other the chain contained 32 consecutive and closely-packed permanent loops of size 30 *σ*. **A.** Average transcriptional activity for each TU in the looped set (magenta bars indicate values for TUs in loops, and magenta arcs loop positions). **B.** Comparison between expression in wild-type and looped configuration for 31 TUs with significantly different values in the two sets (*p* ≃ 0.003; Students t-test). **C.** Regulatory network inferred from the matrix of Pearson correlations between expression of TUs (as Fig. 3A). **D.** Change in Pearson correlation between TUs due to introduction of loops.

**FIG. S4:**
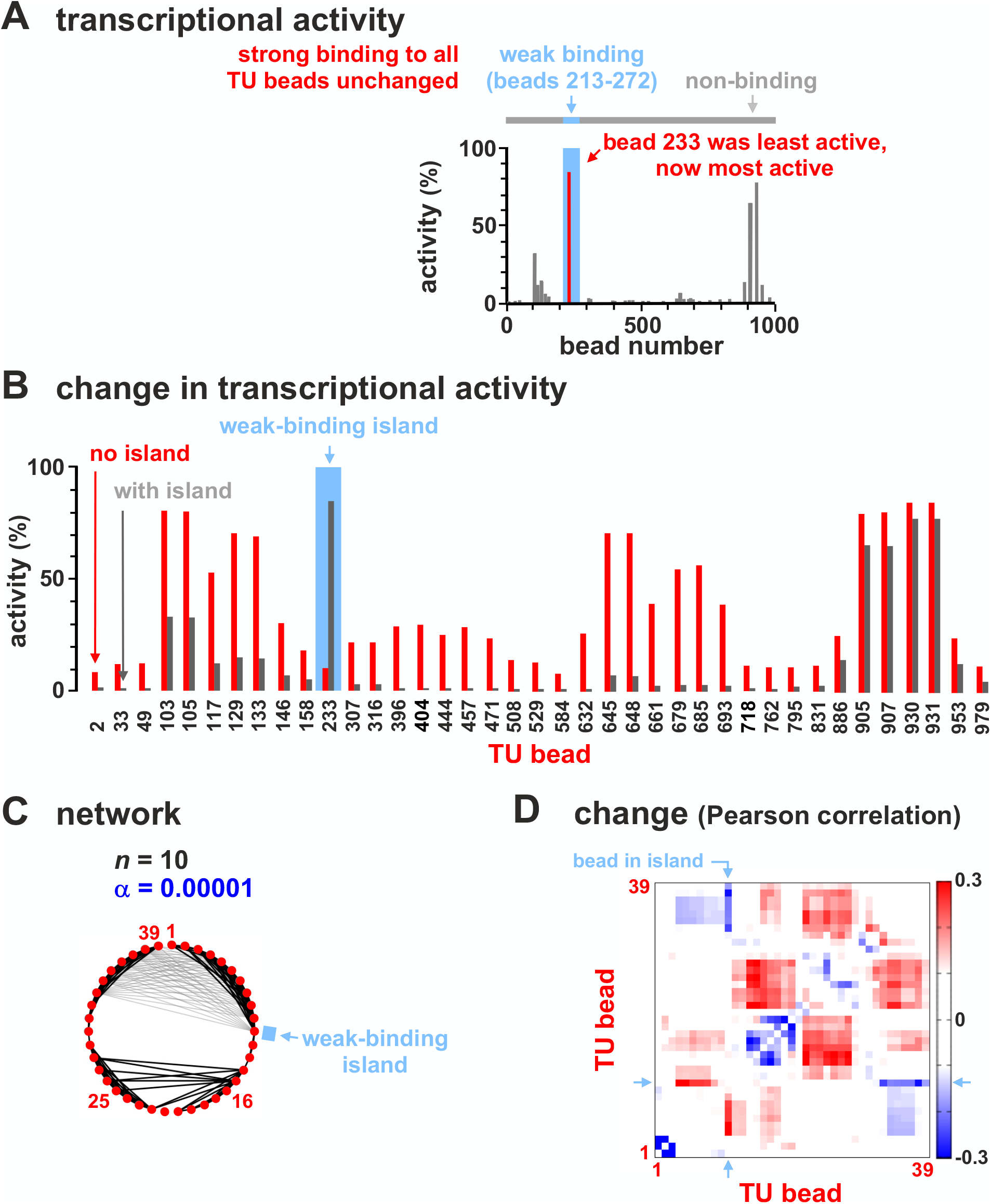
Transcriptional activity in a heterochromatic chain with an euchromatic island. Results from two sets of simulations (800 simulations/condition) were compared; one set was as Fig. 1, in the other all non-TU beads were heterochromatic apart from those between bead 213 and 272 inclusive (to represent a euchromatic island). **A.** Average transcriptional activity for each TU bead. Grey bars: values for TUs in heterochromatin. **B.** Comparison of average transcriptional activity with respect to the wild-type for all 39 TUs; these all have significantly-different values in the two sets (*p* ≃ 0.003; Students t-test). **C.** Regulatory network inferred from the matrix of Pearson correlations between transcriptional activities of TUs (as Fig. 3A). **D.** Change in Pearson correlation between TUs with respect to the wild-type.

**FIG. S5:**
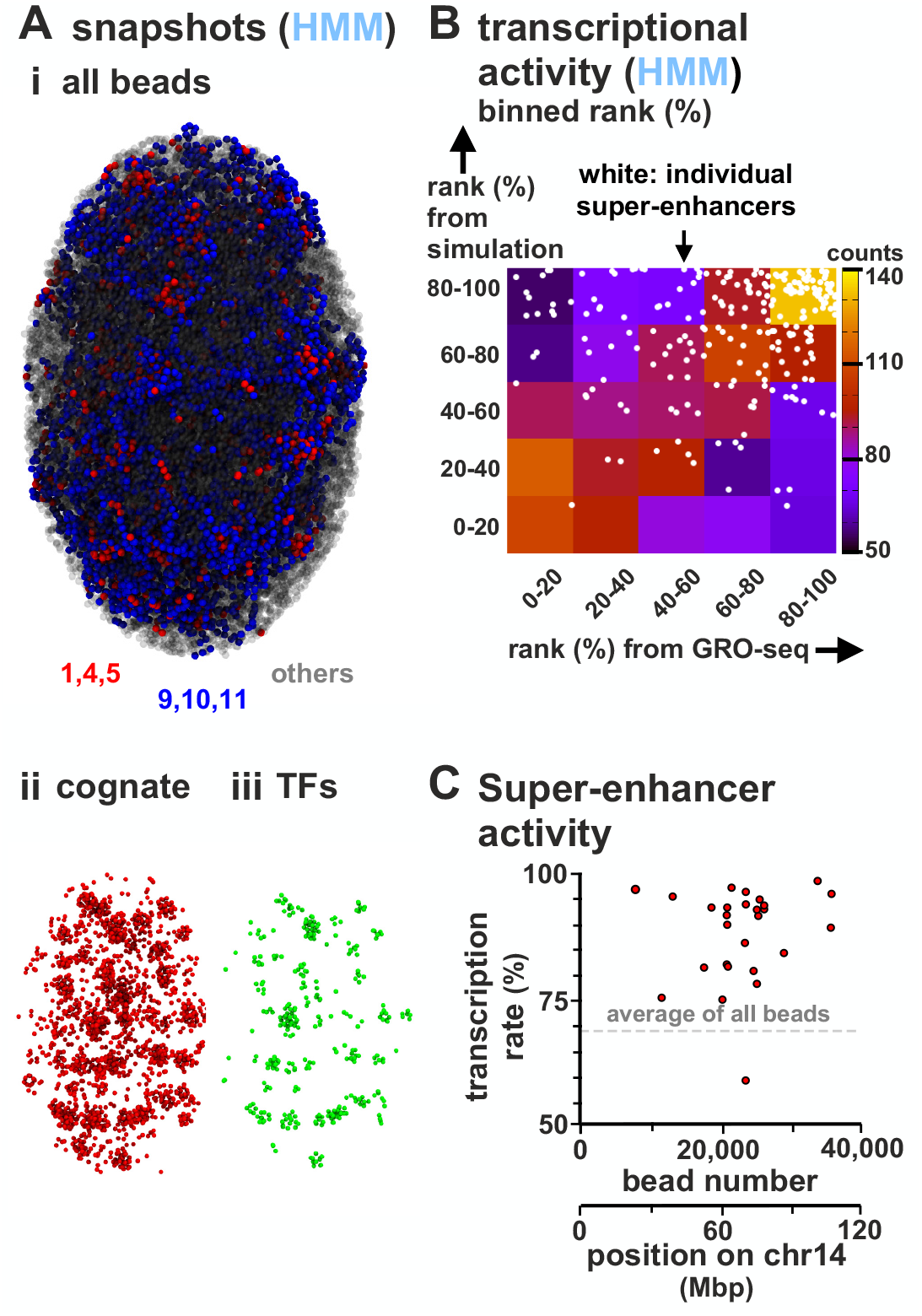
Comparison of transcriptional activities of human chromosome 14 in HUVECs determined using simulations (HMM model; 100 simulations) and GRO-seq. **A. Snapshot.** (i) All beads on the chain (TFs not shown). (ii, iii) TU beads and TFs corresponding to the configuration in (i). **B.** Comparison of transcriptional activities of red TUs in simulations and GRO-seq (ranked from 0 − 100% and binned in quintiles). A scatter plot of unbinned ranks of beads corresponding to SEs are superimposed (white circles). **C.** Transcription rate of SEs. For a given SE, the rate is the average of all TUs in the SE region. We find 26 of 27 SEs have a higher-than-average expression/transcriptional activity.

**FIG. S6:**
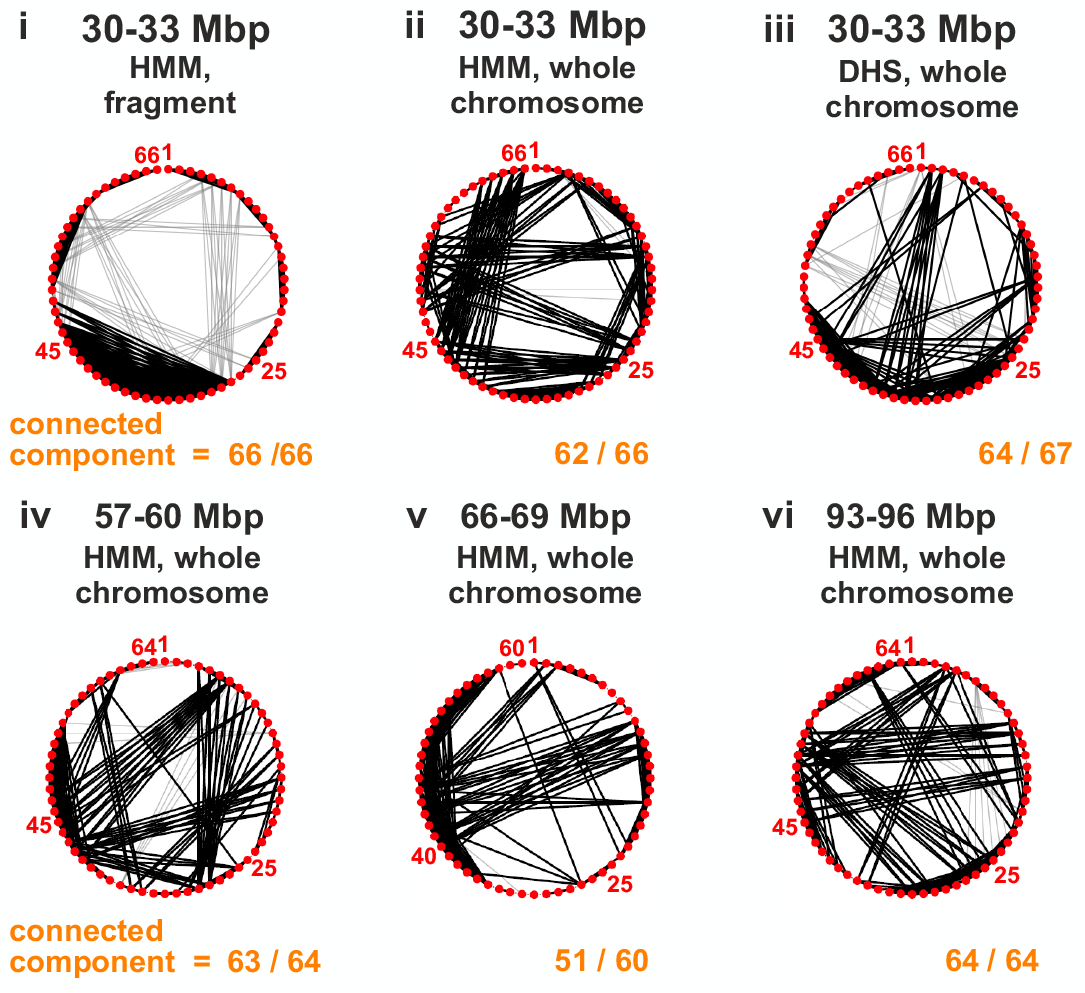
Regulatory networks in chains representing fragments of human chromosome 14 in HUVECs are highly connected and small-world. Networks were constructed and presented as for Fig. 3; they involve Pearson correlations between transcriptional activities of all TUs/red beads. To facilitate comparison with results in Fig. 3, four fragments of the chain representing chromosome 14 were selected that had the same length as the 3 Mbp chain (Fig. 3), and roughly the same density of TUs. (i) Dilute conditions (volume fraction < 0.1%). Simulations (800 runs) involved a short 3 Mbp chain in a cube under the dilute conditions used for the 3 Mbp chain (as Fig. 3), but with beads coloured using the HMM model (as Fig. 7). Consequently, the statistical certainty associated with this panel is inevitably higher than in the other panels as ≥ 8-fold more runs were involved. (ii-vi) Confined conditions (volume fraction ~ 14%). Simulations (100 runs) were conducted using the DHS and HMM models, and a string representing the whole chromosome in an ellipsoid (as Fig. 7, Fig. S5). Networks in all panels are highly connected. Comparison of panels (i) and (ii) – which allow comparison of the effects of confinement – points to confinement increasing the number of distant positive correlations (as might be expected). Comparison of panels (ii) and (iii) which allow comparison of the HMM and DHS models highlights the different patterns of marks contained within the two models. Comparison of panels (ii)-(vi) shows different segments of the chromosome have highly-connected components.

**FIG. S7:**
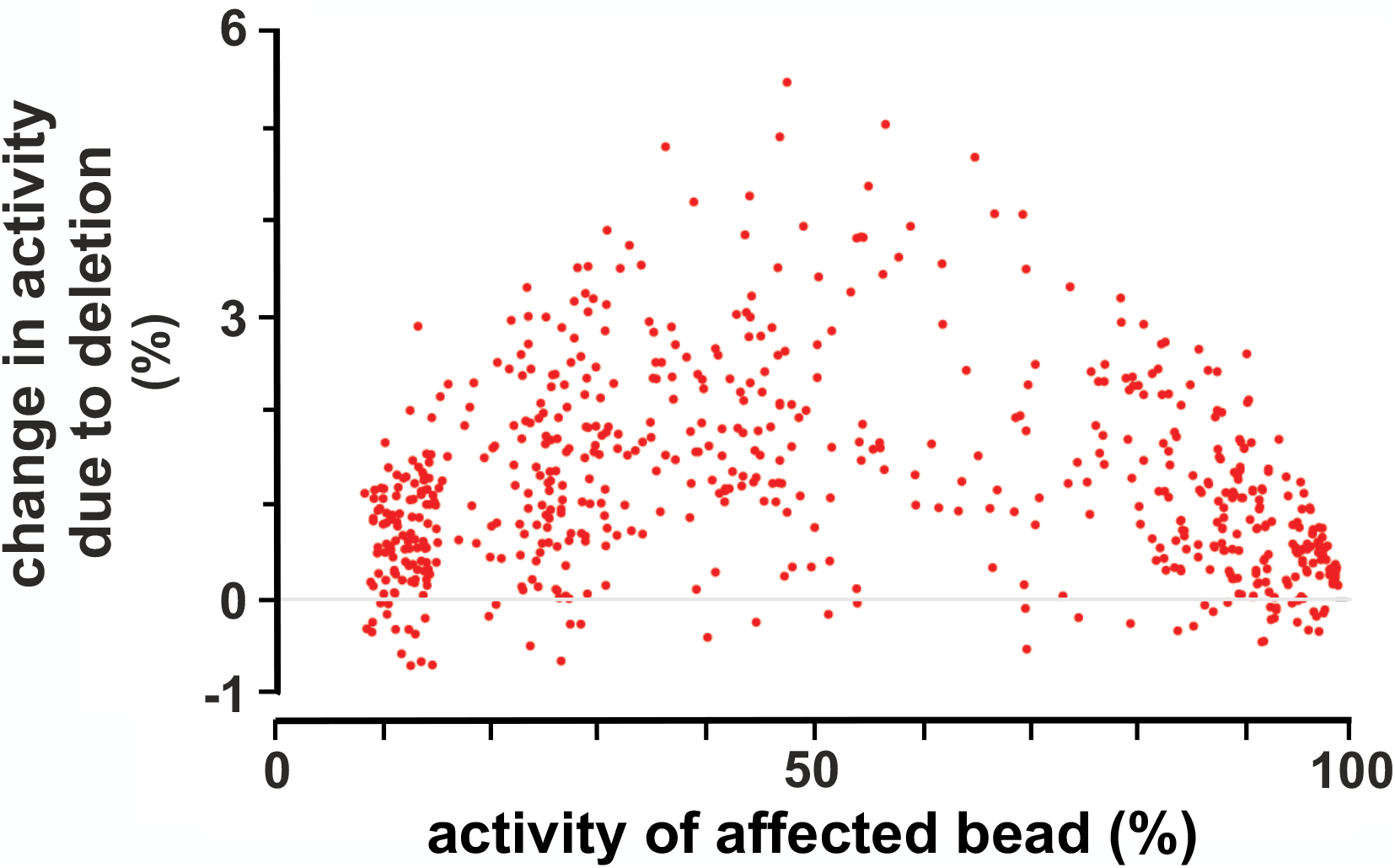
Change in transcriptional activity versus TU activity for DiGeorge deletion. For each TU, we computed both the change in transcriptional activity in the chromosome 22 deletion studied in Fig. 8, as well as the average transcriptional activity in the wild-type and deletion. This scatter plot shows that the change is larger for TUs with intermediate activity.

**FIG. S8:**
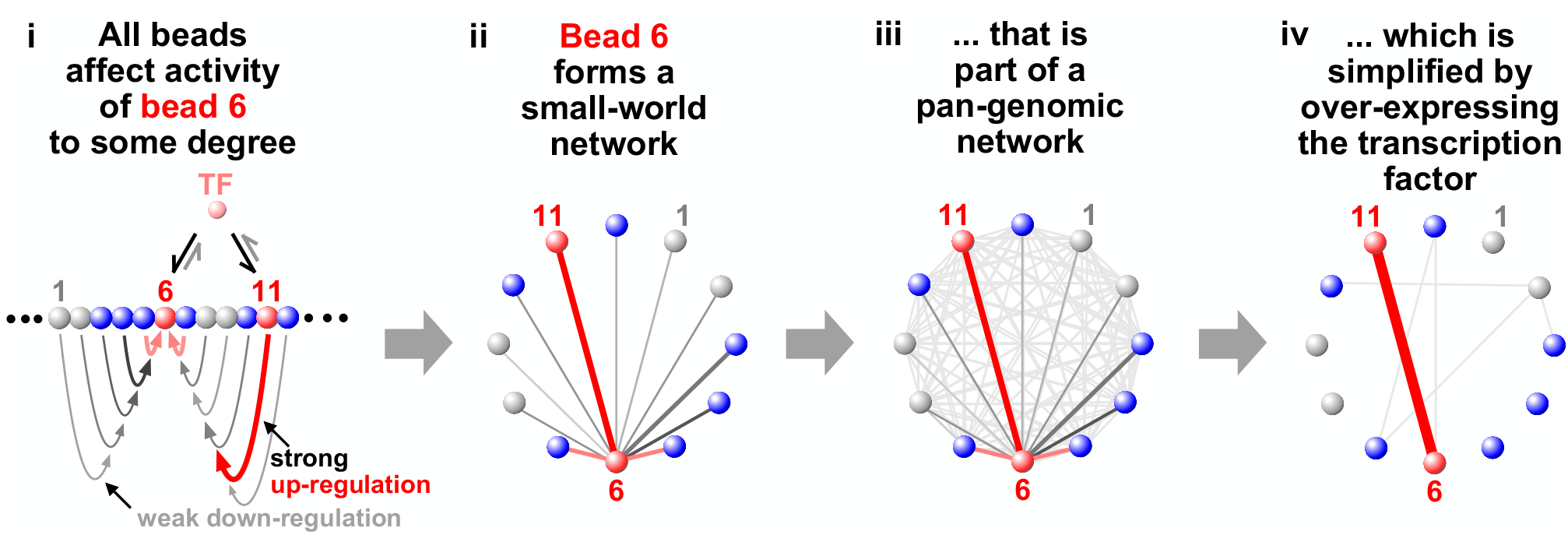
A pan-genomic model for transcriptional organisation and gene regulation. (i) A region of the human genome is depicted with 12 segments (grey – heterochromatin, blue – open chromatin, red – transcription unit). TFs (transcription factor – polymerizing complexes) are present at non-saturating concentrations; they bind tightly to TUs *6* and *11*, weakly to blue segments, and not to heterochromatin. All other beads in the chain influence transcription of bead *6*. They can be considered as eQTLs acting on bead *6*, with eQTL strength being indicated by curved arrows (red – up-regulation, grey – down-regulation; increasing width/colour indicates increasing strength). (ii) The regulatory network centred just on bead 6. Straight lines indicate interaction strength (colour code as i). (iii) The regulatory network of all 12 beads. Every bead has some effect on the transcription of every other bead, which is consistent with GWAS results. (iv) Increasing TF copy-number (as in reprogramming experiments [4]) simplifies both the network of bead *6*, and the complete network. Consequently, over-expressing the TF over-rides the regulatory network mediated by 3D structure, and allows the coexisting trans-acting biochemical network envisioned by the omnigenic model to dominate.

